# Identification of an Arabidopsis Aminotransferase that Facilitates Tryptophan and Auxin Homeostasis

**DOI:** 10.1101/013821

**Authors:** Michael Pieck, Youxi Yuan, Jason Godfrey, Christopher Fisher, Sanda Zolj, Nicholas Thomas, Connie Wu, Julian Ramos, Norman Lee, Jennifer Normanly, John Celenza

**Affiliations:** Department of Biology, Boston University, Boston, MA; Department of Biochemistry and Molecular Biology, University of Massachusetts, Amherst, MA; Chemistry Instrumentation Center, Boston University, Boston, MA

**Keywords:** *Arabidopsis thaliana*, Auxin, ISS1/VAS1, phenylpropanoids, tryptophan metabolism

## Abstract

IAA plays a critical role in regulating numerous aspects of plant growth and development. While there is much genetic support for tryptophan-dependent (Trp-D) IAA synthesis pathways, there is little genetic evidence for tryptophan-independent (Trp-I) IAA synthesis pathways. Using Arabidopsis, we identified two mutant alleles of *ISS1* (*<u>I</u>ndole <u>S</u>evere <u>S</u>ensitive*) that display indole-dependent IAA overproduction phenotypes including leaf epinasty and adventitious rooting. Stable isotope labeling showed that *iss1*, but not WT, uses primarily Trp-I IAA synthesis when grown on indole-supplemented medium. In contrast, both *iss1* and WT use primarily Trp-D IAA synthesis when grown on unsupplemented medium. *iss1* seedlings produce 8-fold higher levels of IAA when grown on indole and surprisingly have a 174-fold increase in Trp. These findings indicate that the *iss1* mutant’s increase in Trp-I IAA synthesis is due to a loss of Trp catabolism. *ISS1* was identified as At1g80360, a predicted aromatic aminotransferase, and in vitro and in vivo analysis confirmed this activity. At1g80360 was previously shown to primarily carry out the conversion of indole-3-pyruvic acid to Trp as an IAA homeostatic mechanism in young seedlings. Our results suggest that in addition to this activity, in more mature plants *ISS1* has a role in Trp catabolism and possibly in the metabolism of other aromatic amino acids. We postulate that this loss of Trp catabolism impacts the use of Trp-D and/or Trp-I IAA synthesis pathways.

## Introduction

In plants, the aromatic amino acids tryptophan (Trp), tyrosine (Tyr) and phenylalanine (Phe) are used for the synthesis of proteins and as precursors to a variety of specialized metabolites. While most secondary metabolites help protect the plant against abiotic and biotic stress (Tzin and Galili 2010), some such as Trp-derived indole-3-acetic acid (IAA) are essential growth regulators (Woodward and Bartel 2005). IAA, the primary auxin in plants, functions in establishing cell polarity during embryogenesis (Weijers et al. 2005; Kleine-Vehn *et al*. 2008), determination of leaf patterning (Bainbridge *et al*. 2008), and initiation of lateral roots and shoots (Celenza *et al*. 1995; Peret *et al*. 2009).

There are two general routes proposed for IAA biosynthesis in plants: through the Trp-D pathway, or through an indolic precursor of Trp [Trp-independent (Trp-I) pathway] (Woodward and Bartel 2005)(Figure 1). Within the Trp-D IAA biosynthetic pathway, there are three established pathways in plants, which lead to IAA production: (i) the indole-3-pyruvic acid (IPA) pathway (ii) the indole-3-acetaldoxime (IAOx) pathway, and (iii) the indole-3-acetamide (IAM) pathway (Figure 1) (Ljung 2013; Zhao 2014).

**Figure 1.**
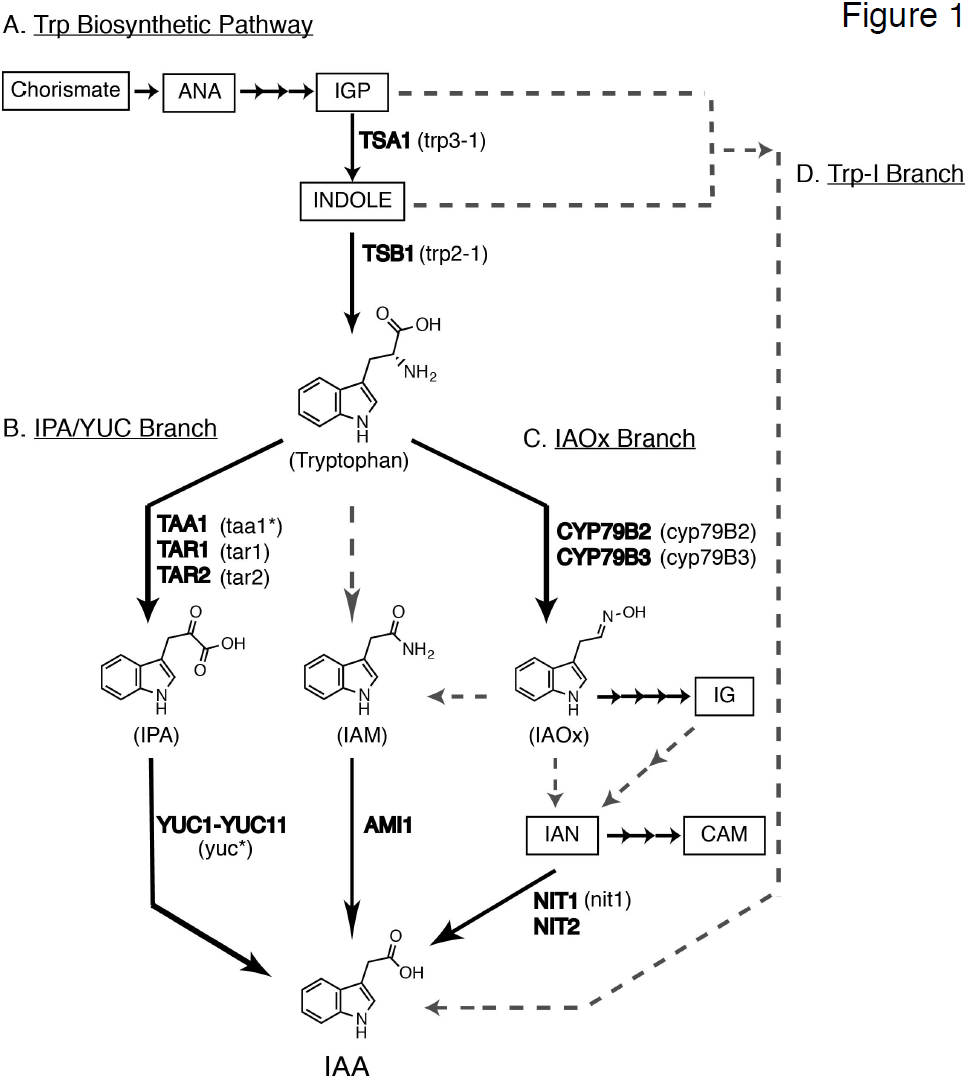
IAA biosynthetic pathway in Arabidopsis. (A) Simplified Trp biosynthetic pathway; IGP, indole-3-glycerolphosphate; *TSA1,* tryptophan synthase alpha; *TSB1,* tryptophan synthase beta. (B) IPA/YUC Branch; *TAA1* <u>Tr</u>yptophan <u>A</u>minotransferase of <u>A</u>rabidopsis-1; *TAR1*; (*taal\**) denotes multiple mutant alleles: *sav3*, *wei8*, *tir2, ckrc1 <u>TA</u>A1-<u>R</u>elated-1 and <u>TA</u>A1-<u>R</u>elated-2;* IPA, indole-3-pyruvic acid; *YUC1*-*YUC11, YUCCA1*-*YUCCA11*; (*yuc\**) denotes multiple yuc mutants: *yuc1, yuc1-D, yuc2, yuc3, yuc4, yuc5, yuc6, yuc6*-*1D, yuc6*-*2D, yuc7, yuc8, yuc9, yuc10*. (C) IAOx Branch; *CYP79B2*, cytochrome P450 (79B2); *CYP79B3*, cytochrome P450 (79B3); IAOx, indole-3-acetaldoxime; IG, indole-3-glucosinolate; CAM, camalexin; IAN, indole-3-acetonitrile; *NIT1*-*3*, nitrilase1 -3; IAM, indole-3-acetamide; *AMI1*, indole-3-acetamide hydrolase-1. (D) Trp-independent (Trp-I) Branch with IGP or indole as the starting substrate. Dashed lines indicate the gene or enzyme has not been identified.

In the IPA pathway, the TAA1 family of Trp aminotransferases converts Trp into IPA (Stepanova *et al*. 2008; Tao *et al*. 2008; Yamada *et al*. 2009; Zhou *et al*. 2011) followed by the direct conversion to IAA by the *YUCCA* (*YUC*) family of flavin monooxygenase-like (FMO) enzymes (Mashiguchi *et al*. 2011; Won *et al*. 2011; Zhao 2012). Both *TAA1* and *YUC* gene families are highly conserved across the plant kingdom and *TAA1* and *YUC* genes are co-expressed in a spatial and temporal manner (Stepanova *et al*. 2011) consistent with TAA1 and YUC being the primary route for IAA biosynthesis.

In Arabidopsis, IAOx is produced by CYP79B2 and CYP79B3 and is an intermediate in the synthesis of IAA and two classes of defense compounds, indole-glucosinolates (IGs) and camalexin (Hull *et al*. 2000; Zhao *et al*. 2002; Glawischnig *et al*. 2004; Sugawara *et al*. 2009). Disruption of the metabolic flux from IAOx to IGs in the *sur1-1* (Mikkelsen *et al*. 2004) or *sur2-1* (Morant *et al*. 2010) mutant causes an increase in conversion of IAOx to IAA, supporting this pathway as a route to IAA. However, IAOx is likely a minor contributor to overall IAA production under most conditions because the double *cyp79B2 cyp79B3* mutant produces no IAOx and appears WT (Zhao *et al*. 2002; Sugawara *et al*. 2009). In addition, IAOx production appears to be limited to the Brassicaceae family (Sugawara *et al*. 2009; Nonhebel *et al*. 2011).

For the IAM pathway, IAM is produced from Trp, perhaps by a monooxygenase, similar to what is used by bacteria (Woodward and Bartel 2005; Ljung 2013), although in Arabidopsis the IAOx pathway may also produce IAM (Sugawara *et al*. 2009). In Arabidopsis IAM can be converted to IAA in vitro by IAM hydrolase encoded by *Amidase 1* (*AMI1*) (Pollmann *et al*. 2003). While the IAM pathway has been proposed to exist throughout the plant kingdom (Mano *et al*. 2010) its importance in overall IAA metabolism remains to be characterized genetically.

Evidence of the Trp-I IAA synthesis pathway comes primarily from stable isotope labeling studies of Trp auxotrophs in maize (Wright *et al*. 1991) and Arabidopsis (Normanly *et al*. 1993), IAA homeostasis mutants (Quint *et al*. 2009) and in wild type *Lemna gibba* under environmental perturbations (Rapparini *et al*. 2002). Indole or indole-3-glycerol phosphate, precursors of Trp, have been proposed as the branch point to Trp-I IAA biosynthesis (Ouyang *et al*. 2000). Indirect evidence comes from analysis of the Arabidopsis *alf3-1* mutant, which produces lateral roots that arrest growth and die after the initial stages of lateral root formation (Celenza *et al*. 1995). However, the *alf3-1* phenotype is rescued by medium supplemented with indole but not with Trp (Celenza *et al*. 1995), suggesting that the rescue is via Trp-I IAA biosynthesis.

In this work we describe the identification of *<u>i</u>ndole <u>s</u>evere <u>s</u>ensitive1* (*iss1*), a mutant that displays indole-dependent adventitious roots, leaf narrowing and longer petioles due to elevated Trp-I IAA synthesis. Indole-grown *iss1* mutants have elevated IAA and Trp levels as well as alterations in phenylpropanoid metabolism. *ISS1* encodes an aromatic aminotransferase (AroAT); an allele of *ISS1*, named *vas1*, was identified previously in a screen for suppressors of *taa1* and was found to carry out the conversion of IPA to Trp using methionine as an amino donor(Zheng *et al*. 2013). The results presented herein provide evidence that ISS1/VAS1 plays a role in Trp catabolism in addition to its role in modulating IPA levels and can function as a more general AroAT.

## Materials and Methods

### Plant materials and growth conditions

All plants listed are in the Columbia (Col-0) background except *iss1-2*, which is in the Wassilewskija (Ws) background. *iss1-1* was identified by screening pools of random T-DNA insertion lines in the Col-0 background [CS76502, CS76504, CS76506 and CS76508; Arabidopsis Biological Resource Center (http://abrc.osu.edu/)] grown on PNS medium containing 80 μM indole (PNS+indole). *iss1-2* was identified by screening EMS-mutagenized Wassilewskija (Ws) seeds on PNS+indole. The *wei8-4* allele of *TAA1* (SALK_022743C) and *trp2-1* (CS8327) were obtained from the ABRC.

For positional cloning, the homozygous *iss1-2* mutant in the Ws background was crossed to WT (Col-0) and the F_1_ progeny were allowed to self-fertilize. The F_2_ progeny were grown on PNS+indole and approximately 1100 plants displaying the *iss1* phenotype were selected as a mapping population. Microsatellite markers were used to isolate the *iss1* locus to a region on chromosome 1 between BAC clones F18B13 and F5I6 using primers listed in (Table S1.)

To generate mutant combinations, strains were crossed and the F1 progeny were allowed to self-fertilize. Putative double or triple mutants were identified in subsequent generations by PCR genotyping (Table S2).

*35S::ISS1* constructs were generated using Univector Plasmid-fusion System (UPS) (Liu *et al*. 1998) using the pKYLX-myc9-loxP host vector (Guo and Ecker 2003). For plant transformation, the floral dip method was used (Clough and Bent 1998) and three homozygous *35S::ISS1* lines were selected by kanamycin resistance.

Surface sterilized seeds were grown on plant nutrient medium containing 0.5% (w/v) sucrose (PNS) and solidified with 0.6% agar, if needed (Haughn and Somerville 1986). Unless otherwise noted, PNS+indole contained 80 μM indole; Trp, 5-methyl-tryptophan (5MT), or p-fluorophenylalanine (PFP) supplements are as indicated. Plants were grown at 22° under constant light using a yellow low pass filter at a light intensity of 20-40 μE m^-2^ s^-2^. For UV-B treatment, plants were incubated at RT for 24 h with a UV-B light intensity of 0.8-1.1 μE m^-2^ s^-2^. Root growth measurements were performed using ImageJ (http://imagej.nih.gov/ij/) and are reported as the mean root length +/- SE with analysis by a two-tailed Student’s t-test.

### Quantification of IAA, Trp, Phe and Tyr and secondary metabolites

Unless otherwise noted, all metabolite measurements are reported as the average of three samples +/- SE and analyzed statistically as indicated in the figure legend. Solid phase extraction of IAA was performed on 30-75 mg of plant tissue according to (Barkawi *et al*. 2010). Methylated IAA samples were subjected to GS-SIM-MS analysis as described by (Barkawi *et al*. 2010). 100 pg/μl of authentic methyl-IAA was run to determine the retention time. The molecular and quinolinium ions of endogenous IAA and the [^13^C_6_]-IAA internal standard (Cambridge Isotope Laboratories, Tewksbury, MA), are listed in Table S3. Levels of free IAA were quantified using the quinolinium ion as described by (Barkawi *et al*. 2010).

Extraction and analysis of Trp, Phe and Tyr was performed according to (Chen *et al*. 2010) using methyl chloroformate derivitization. For Trp and Phe quantification, known amounts of [^2^H_5_]-Trp, and [^2^H_5_]-Phe (Cambridge Isotope Laboratories, Tewksbury, MA) were included as internal standards. Tyr was quantified using a standard curve of unlabeled Tyr. The molecular and major fragment ions of Trp, [^2^H_5_]-Trp, Phe, [^2^H_5_]-Phe, and Tyr are listed in Table S3.

Glucosinolates were isolated from methanol extracts using anion exchange (DEAE Sephadex A25) and converted to desulfoglucosinolates by adding aryl sulfatase as described (Brown *et al*. 2003). HPLC of desulfoglucosinolates was carried out using a Waters 2795 HPLC equipped with Waters 2996 photodiode array detector and a Phenomenex Luna 5 micron 250 × 4.6 mm C18 column. Desulfoglucosinolates were confirmed by comparing the retention time and UV spectra to purified standards and quantified at 229 nm relative to an external sinigrin standard (Brown *et al*. 2003).

Phenylpropanoids were quantified from methanol extracts using a Waters 2795 HPLC equipped with a Waters 2996 photodiode array detector and a Phenomenex Luna 5 micron 250 × 4.6 mm C18 column. The mobile phase was a gradient of water with 0.1% formic acid (A) and acetonitrile with 0.1% formic acid (B) as follows: 95% A; 1-25 min, linear gradient to 40% A; 25-28 min, linear gradient to 100% B; 28-33 min, 100% B; 33-37 min linear gradient to 95% A; 37-42 min 95% A. Detection of separated compounds was monitored using full spectrum detection from 200 nm to 400 nm. Compounds were identified by comparison of RT and UV spectra compared to previously published results (Hemm *et al*. 2004; Kerhoas *et al*. 2006) and quantified using chromatographs extracted at 260 nm relative to WT control. Peak areas were normalized to wet tissue weight. For UV-B inducible flavonoid quantification, the peak areas of the two most abundant UV-B responsive flavonoids were summed.

The mass of coniferin was confirmed by high-resolution mass spectrometry (HRMS) obtained on a Waters Q-ToF (hybrid quadrupole/time-of-flight) API US system (Waters, Milford, MA) by electrospray (ESI) in the [positive] mode. Mass correction was done by an external reference using a Waters Lockspray accessory. Mobile phases were water and acetonitrile with 0.1% formic acid. The MS settings were: capillary voltage = 3kV, cone voltage = 35, source temperature = 120° and dissolvation temperature = 350°.

### Dual stable isotope labeling

Plants were grown on PNS+indole plates until the *iss1-1* phenotype was observed, typically 20 days after germination. 4-6 WT plants or 10-12 mutant plants were transferred to a sterile 6-well tissue culture plate. Labeling was initiated by adding 3 ml of liquid PNS containing 10 μM [^15^N] ANA (Cambridge Isotope Laboratories, Tewksbury, MA) and 10 μM [^13^C_11_ ^15^N_2_] TRP (Cambridge Isotope Laboratories, Tewksbury, MA) in, and returning the plates to the growth chamber for 12 h. (Because [^15^N] ANA and [^13^C_11_ ^15^N_2_] TRP were dissolved in 2-propanol, a mock labeling of 2-propanol alone was done to determine that the addition of the solvent did not perturb IAA synthesis.) Tissue was then transferred to a mesh screen that was stretched over a beaker and rinsed with sterile deionized H2O to remove excess label and media. The tissue was briefly patted dry with paper towels and then weighed, frozen on dry ice and stored at -80°. IAA and Trp extraction and quantification were performed as described above with 5 ng [^13^C_6_]-IAA and 1 μg of [^2^H_5_]-Trp added as internal standards. The quinolinium ion of each isotopomer is listed in Table S3 and was used for isotope enrichment analysis. The percent incorporation of each isotopomer for Trp and each isotopomer for IAA was calculated from the total amount of Trp and IAA, respectively (using the internal standards [^13^C_6_] IAA and [^2^H_5_]Trp).

Using the percent incorporation for each isotopomer, the percent IAA_Trp-D_ and IAA_Trp-I_ were calculated using the following formulas as previously suggested (Liu *et al*. 2012): IAA_Trp-D_ = [^13^C_10_^15^N]IAA + [^15^N]IAA_Trp-D_ and IAA_Trp-I_ = [^15^N]IAA_Trp-I_. Since [^13^C_11_ ^15^N_2_]Trp can only be converted to [^13^C_10_ ^15^N]IAA via the Trp-D pathway, any [^13^C_10_ ^15^N]IAA detected is from the Trp-D pathway. Additionally, any [^15^N]Trp converted to [^15^N]IAA from the Trp-D pathway should be proportional to the conversion of [^13^C_11_ ^15^N_2_]Trp to [^13^C_10_ ^15^N]IAA. Therefore, the ratio of Trp-dependent [^15^N]IAA to [^13^C_10_^15^N]IAA is equal to the ratio of [^15^N]Trp to [^13^C_11_ ^15^N_2_]Trp. Or [^15^N]-IAA_Trp-D_ = [^13^C_10_ ^15^N]IAA x [^15^N]Trp / [^13^C_11_ ^15^N_2_]Trp. Once the amount of [^15^N]IAA_Trp-D_ was calculated, the amount of [^15^N]-IAA_Trp-I_ can be calculated by subtracting [^15^N]IAA_Trp-D_ from [^15^N]IAA_total_. Or [^15^N]IAA_Trp-I_ = [^15^N]IAA_total_ - [^15^N]IAA_Trp-D_.

### Heterologous expression constructs

For the expression of Arabidopsis cDNAs in yeast and plants, the Univector Plasmid-fusion System (UPS) was used (Liu *et al*. 1998). UPS-compatible Arabidopsis cDNAs and expression vectors (pHOST) are listed in Table S4. The *iss1-2* mutation was introduced into *ISS1* cDNA carried on an *E. coli* plasmid by using PCR and Gibson assembly according to supplier’s instructions (New England Biolabs, Ipswich, MA).

### Complementation of the yeast aro8 aro9 Phe and Tyr auxotrophy

Yeast strains 11965 and 14569 in the S288C background (GE Dharmacon, Lafayette, CO) were used to generate a haploid *aro8::HygR aro9::G418R* double mutant (MPY1; Table S5) for complementation assays. *ISS1, iss1-2, TAA1* and *MEE17* cDNAs recombined into pHY326-loxH were transformed into MPY1 and grown on yeast minimal (SD) medium (Ausubel *et al*. 1994-1998), with or without 50 mg/l Phe, and 30 mg/l Tyr, and supplemented with either 2% glucose or 2% galactose. Complementation was assayed following incubation at 30° for 48 h.

For pseudohyphal growth assays, diploid yeast strains PY652 and PY653 (∑1278B background) were obtained from Dr. Prusty-Rao (Worcester Polytechnic Institute, Worcester, MA) and strain MPY5 was derived from PY653 (Table S5). To induce pseudohyphal growth, yeast was grown on solid synthetic low ammonium dextrose (SLAD) medium with or without Phe and Tyr supplemented with either 0.5% glucose or 0.5% galactose. Pseudohyphal growth was observed following incubation at 30° for 72-96 h.

### HPLC based aromatic aminotransferase enzyme assay

*ISS1* and *iss1*-*2* cDNAs were subcloned into the IPTG inducible pMAL-c4X MBP expression vector (New England Biolabs, Ipswich, MA). MBP fusion protein was purified according to the supplier’s instructions (New England Biolabs, Ipswich, MA). Purified protein was quantified by Bradford assay. The presence of recombinant protein was confirmed by SDS-PAGE and native PAGE analysis. An empty pMAL-c4X vector was used as a control.

AroAT activity was measured using a method adapted from (Stepanova *et al*. 2008) as follows. The reaction buffer consisted of 0.1 M borate pH 6.5, 4 mM EDTA, 0.2 mM PLP, 5 mM amino donor and 0.5 mM amino acceptor. The reaction was initiated by adding 5-60 μg/ 200 μL purified MBP-tagged protein. A zero-time point was taken immediately after adding the protein. The reaction was incubated at 37° for 30, 60 and 120 min. At each time point, 200 μL was removed from the reaction volume and added to a 1.5 ml microcentrifuge tube containing 200 μL methanol to stop the reaction. The stopped reaction was briefly vortexed and centrifuged for 30 sec. After centrifugation, 100 μL was analyzed for production of the aromatic amino acid aromatic α-keto acid were analyzed using a Waters 2795 HPLC equipped with a Waters 2996 photodiode array detector and a Phenomenex Luna 5 micron 250 × 4.6 mm C18 column. The mobile phase was a gradient of water with 0.1% formic acid (A) and acetonitrile with 0.1% formic acid (B) as follows: 90% A; 1-5 min; 5-10 min, linear gradient to 85% A; 10-20 min, linear gradient to 83% A; 20-30 min, linear gradient to 75% A; 30-40 min, linear gradient to 65% A; 40-50 min, linear gradient to 55% A; 55-58 min, linear gradient to 45% A; 58-60 min, linear gradient to 10% A; 61-66 min 90% A. Detection of separated compounds was monitored at 254 nm and 378nm. Products were compared to the RT of authentic standards for Trp, Phe, Tyr, indole-3-pyruvate, phenylpyruvate, and 4-hydroxyphenylpyruvate.

## Results

### The *iss1*-*1* mutant displays an indole-dependent high auxin phenotype

Because the branch point between Trp-I IAA biosynthesis and Trp-D IAA biosynthesis is proposed to be an indolic precursor to Trp (Woodward and Bartel 2005), we reasoned that mutants displaying a high-IAA phenotype when supplemented with indole, but not Trp, would have an increase in Trp-I IAA biosynthesis. We screened through a publicly available single T-DNA insertion collection from which one seedling, *<u>i</u>ndole <u>s</u>evere <u>s</u>ensitive* (*iss1*-*1*) developed a high IAA phenotype that was observable after 2-3 weeks growth on Plant Nutrient Sucrose (PNS) medium supplemented with 80 mM indole (PNS+indole). Compared to the wild-type Columbia plants, the *iss1*-*1* mutant displayed increased root growth inhibition by indole (Figure 2A-B) and phenotypes consistent with increased auxin including increased lateral root formation, narrow leaves, and elongated petioles (Figure 3). *iss1-1* appeared normal when grown on PNS medium or PNS supplemented with 80 mM Trp (data not shown). After backcrossing to Col-0 the indole sensitivity segregated as a single recessive Mendelian trait. The *iss1*-*2* allele was recovered from an EMS-mutagenized M2 population in the Ws accession, had a similar phenotype to *iss1*-*1*, and failed to complement *iss1*-*1*.

**Figure 2.**
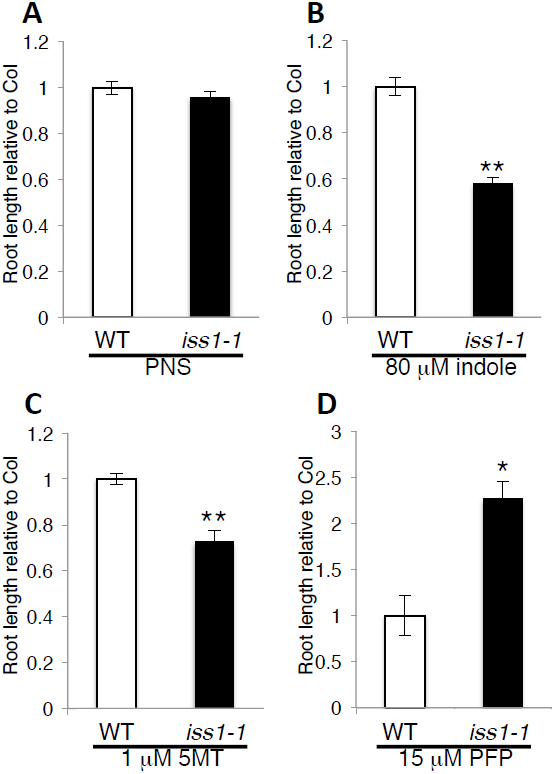
The *iss1*-*1* mutant displays altered root growth sensitivity to indole, 5-methyl tryptophan, and p-fluorophenylalanine. 15-day-old average root length normalized to WT is shown for WT (white bars) and *iss1*-*1* (black bars) plants grown on unsupplemented PNS (A) or PNS+indole (B), PNS + 1 μ5MT (C) or PNS 15μM PFP (D). *iss1*-*1* root growth is significantly different from WT for B, C [P< 0.001 (**) two-tailed Student’s *t*-test] and D [P <0.005 (*) two-tailed Student’s *t*-test].

**Figure 3.**
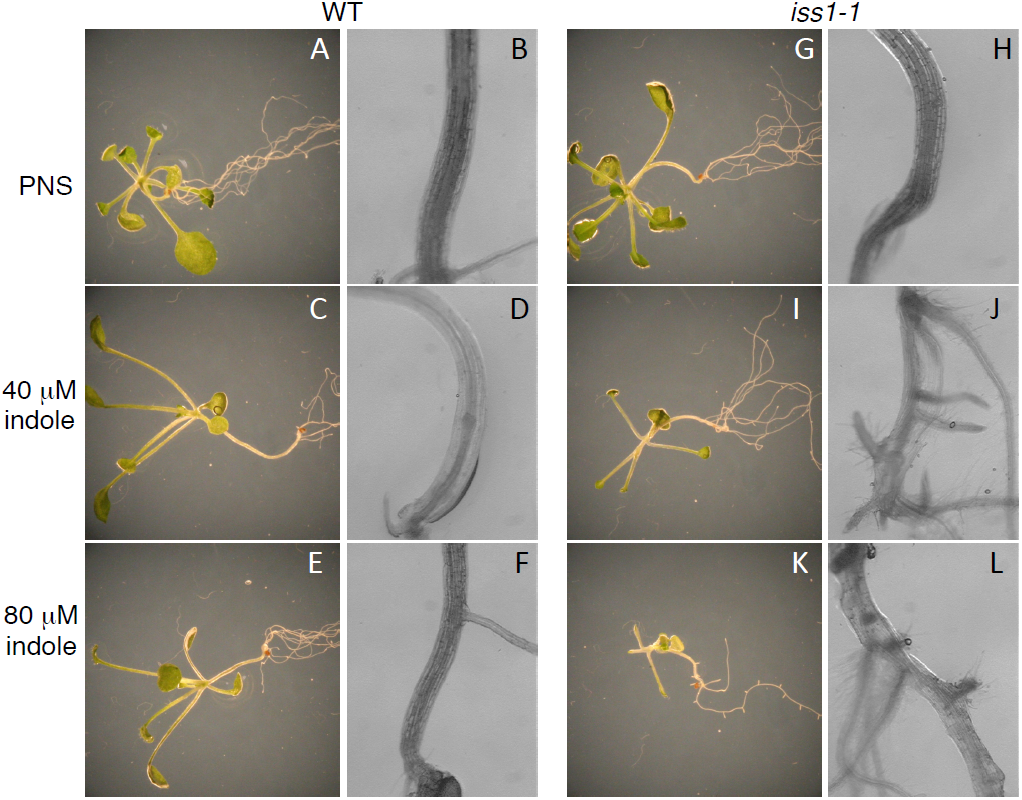
The *iss1*-*1* mutant displays an indole-dependent high IAA phenotype. Depicted are representative 18-day-old WT and *iss1*-*1* mutant plants grown on PNS (A,B, G, H), or PNS containing 40 μM indole (C, D, I, J), or 80 μM indole (E, F, K, L).Shown are dark field images of the seedling (A, C, E, G, I, K) and bright field images of the hypocotyl (B, D, F, H, J, L).

### The *iss1*-*1* mutant has an indole-dependent increase in IAA due to elevated Trp-I IAA biosynthesis

To determine if the high auxin phenotype observed in *iss1*-*1* is indeed caused by an increase in IAA levels, we quantified IAA using gas chromatography-selected ion monitoring-mass spectrometry (GC-SIM-MS). On PNS medium, IAA levels were similar between *iss1*-*1* and WT (Table 1). However, on PNS+indole medium, *iss1*-*1* accumulated ~8-fold greater IAA than WT, consistent with the high IAA phenotype (Table 1).

**Table 1.** *iss1*-*1* mutants have an indole-dependent increase in free IAA levels. Shown are free IAA levels extracted from 21 day old WT and *iss1*-*1* plants grown on PNS or PNS+indole. Fold change is given relative to WT grown on PNS. * indicates P<0.0001 using a two-tailed Student’s *t*-test. Other fold changes are not significant.

| Genotype/treatment | IAA ng/g TW | +/- SE | Fold Change |
| --- | --- | --- | --- |
| WT/PNS | 18.9 | 1.7 | — |
| WT/PNS+indole | 31.3 | 6.9 | 1.6 |
| *iss1-1*/PNS | 24.7 | 3.8 | 1.3 |
| *iss1-1*/PNS+indole | 157.1 | 5.1 | 8.3* |

Because the *iss1*-*1* mutant displayed phenotypes consistent with elevated levels of IAA when grown on indole but not when grown on Trp, we hypothesized that the increase in IAA was due to an increase in Trp-I IAA biosynthesis relative to Trp-D IAA biosynthesis. To distinguish between Trp-D and Trp-I IAA biosynthesis we fed twenty-day old *iss1*-*1* and WT plants grown on either PNS or PNS+indole media the stable isotopes [^15^N] anthranilate (ANA) and [^13^C_11_ ^15^N_2_] Trp for 12 h. The rationale for dual labeling is that any [^13^C_10_ ^15^N] detected in IAA results only from the conversion of [^13^C_11_^15^N_2_] Trp via a Trp-D IAA pathway. In contrast, [^15^N] IAA derives from either [^15^N] ANA first being converted into [^15^N] Trp and then into [^15^N] IAA via a Trp-D IAA pathway or by the conversion of [^15^N] ANA into [^15^N] IAA via a Trp-I IAA pathway (and thus bypassing Trp). Therefore by quantifying the relative conversion of [^13^C_11_^15^N_2_] Trp to [^13^C_10_ ^15^N] IAA and comparing this to the relative amounts of [^15^N] Trp and [^15^N] IAA, the percentage of labeled IAA coming from the Trp-D IAA and Trp-I IAA can be determined (Quint *et al*. 2009; Liu *et al*. 2012).

We first examined relative isotopic enrichment of the Trp pool and found that both *iss1*-*1* and WT grown on PNS or PNS+indole media had similar ratios of [^15^N] Trp to [^13^C_11_ ^15^N_2_] Trp indicating that the uptake of [^15^N] ANA relative to [^13^C_11_ ^15^N_2_] Trp is not changed by genotype or growth condition (Figure 4A). We also examined relative isotopic enrichment of the IAA pool and found a significant increase in the ratio of [^15^N] IAA to [^13^C_10_ ^15^N] IAA for *iss1*-*1* plants grown on PNS+indole medium compared to *iss1-1* grown on PNS medium or WT grown in either condition (Figure 4A). By calculating the increased enrichment of [^15^N] into to IAA relative to Trp, the amount of IAA derived from Trp-I IAA synthesis can be calculated. On PNS, Trp-I IAA synthesis accounted for 38% of the IAA produced in WT and 15% of the IAA produced in *iss1*-*1* (Figure 4B). On PNS+indole medium, the amount of IAA produced via Trp-I IAA synthesis decreased to 18% for WT seedlings, while indole-grown *iss1*-*1* had a dramatic increase to 75% percent (Figure 4B), suggesting that the high levels of IAA in indole-grown *iss1*-*1* derive primarily from the Trp-I IAA pathway.

**Figure 4.**
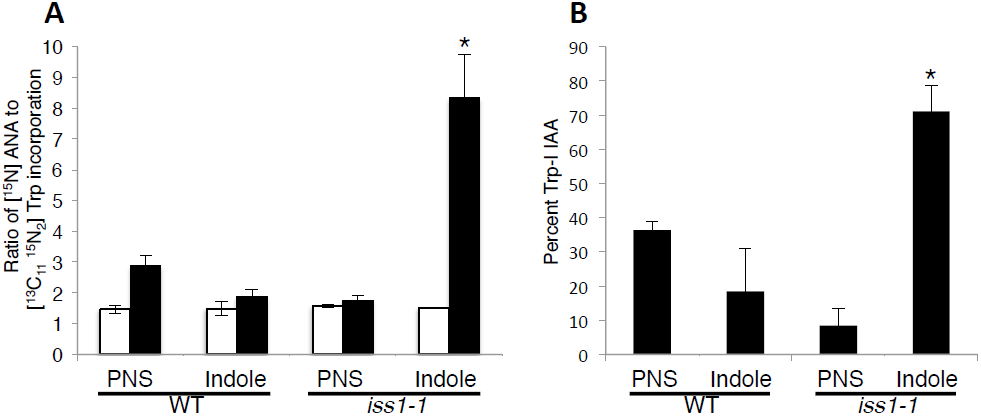
*iss1*-*1* has an indole-dependent increase in Trp-independent IAAbiosynthesis. (A) The ratio of [^15^N] ANA to [^13^C_11_ ^15^N_2_] Trp incorporation into Trp (white bars) and IAA (black bars) is shown. Only the ratio of incorporation into IAA for indolegrown *iss1*-*1* seedlings issignificantly different from other samples (*P < 0.01 two-taile Student’s*t*-test). Details of the labeling procedure are described in Materials and Methods. (B) For the samples used in (A), the percentage IAA made through a Trp-I pathway was determined by calculating the increased enrichment of [^15^N] into to IAA relative to Trp as desciibed in Materials and Methods. Only indole-grown *iss1*-*1* seedlings showed a significant difference in Trp-I IAA synthesis (*P< 0.01 two-tailed Student’s *t*-test).

We also examined IAA synthesis pathway use genetically. Both the IPA and IAOx Trp-D IAA pathways can be reduced or eliminated, respectively, by mutations in key steps in each pathway ((Zhao *et al*. 2002; Stepanova *et al*. 2008; Tao *et al*. 2008). Thus we asked if the *taa1* mutation or the *cyp79B2 cyp79B3* double mutant could suppress *iss1*-*1* indole sensitivity. As seen in Figure 5, the *iss1*-*1 taa1* double mutant and the *iss1*-*1 cyp79B2 cyp79B3* triple mutant remained sensitive to indole.

**Figure 5.**
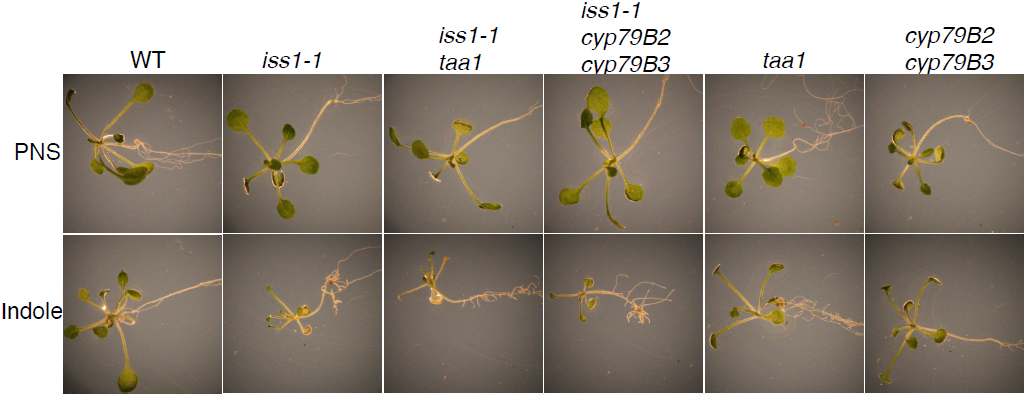
Mutations in Trp-dependent IAA synthesis pathways do not suppress the *iss1*-*1* indole sensitivity. Shown are representative 18-day-old plants of the genotypes shown grown on PNS (top row) or PNS+indole (bottom row).

### The *iss1*-*1* mutant has altered Trp metabolism

Surprisingly, *iss1*-*1* grown on PNS+indole medium also accumulated 174-fold higher levels of Trp suggesting an overall disruption of Trp metabolism (Figure 6). In contrast, phenylalanine (Phe) and tyrosine (Tyr) levels were similar between WT and *iss1-1* plants when grown on PNS or PNS+indole media. Independent of the genotype, we found that growth on PNS+indole relative to PNS medium caused a two-fold increase in Phe and four-fold increase in Tyr (Figure 6).

**Figure 6.**
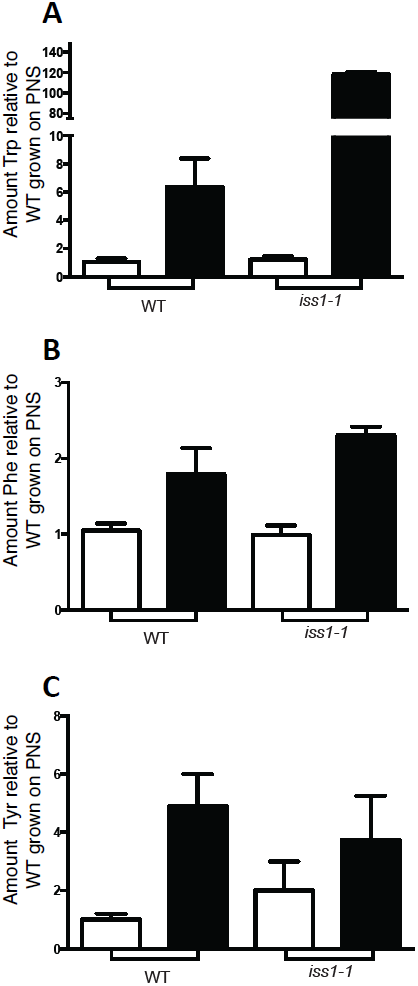
The *iss1*-*1* mutant has altered Trp metabolism. Relative levels of Trp (A),Phe (B) and Tyr (C) are presented normalized to WT plants grown on PNS. Samples were extracted from whole 15-day-old Arabidopsis seedlings grown on PNS (whitebars)or PNS+indole (black bars) and metabolites were detected by GC-MS following derivitization as described in Materials and Methods. The Trp levels in WT grown on PNS+indole are significantly different from WT plants grown on PNS (P< 0.005 using a two-tailed Student’s *t*-test). Trp levels in *iss1*-*1* grown on PNS+indole are significantly different than other samples (P<0.0001 using two-tailed Student’s t-test). Both WT and *iss1*-*1* grown on PNS+indole had higher levels of Phe (P < 0.0005 two-way ANOVA) and Tyr (P <0.005 two-way ANOVA) compared to WT grown on PNS.

Because the *iss1*-*1* mutant accumulated Trp significantly when grown on PNS+indole medium, we predicted that Trp catabolism is altered in the *iss1*-*1* mutant. Therefore we compared *iss1*-*1* growth to WT in the presence of the toxic Trp analog 5MT. Changes in 5MT sensitivity reflect changes in Trp metabolism, such that mutants with decreased Trp catabolism show increased 5MT sensitivity compared to WT (Zhao *et al*. 2002; Celenza *et al*. 2005; Tao *et al*. 2008). The *iss1*-*1* mutant had an increased sensitivity to 5MT compared to WT consistent with a decrease in Trp catabolism (Figure 2C).

### The *trp2* mutant partially suppresses the *iss1*-*1* phenotype

We hypothesized that *iss1*-*1* has altered Trp metabolism, as evidenced by both elevated Trp and IAA levels when *iss1*-*1* was grown in the presence of indole and its increased sensitivity to 5MT. Therefore, to test if Trp synthesis was required for the indole-dependent *iss1*-*1* high IAA phenotype we used double mutant analysis to examine the interaction between *iss1*-*1* and the Trp synthase β (*TSB1*) mutant *trp2*-*1*. The *trp2*-*1* mutant is unable to convert indole to Trp. *trp2*-*1* accumulates indole (Last *et al*. 1991) and exhibits elevated Trp-I IAA biosynthesis (Normanly *et al*. 1993). We found that *trp2*-*1* (Figure 7) suppressed the indole sensitivity phenotype in *iss1*-*1*. In particular, unlike the single *iss1*-*1* mutant, *iss1*-*1 trp2*-*1* plants had normal roots and leaves when grown on PNS+indole medium suggesting that the ability to convert added indole to Trp by TSB1 is needed for the *iss1*-*1* high IAA phenotype. Consistent with this observation, we found that growth of *iss1*-*1 trp2*-*1* on PNS+indole+Trp medium restored the high IAA phenotype. If increased indole alone was responsible for the indole sensitivity of the *iss1*-*1* mutant, then the *iss1*-*1 trp2*-*1* double mutant should have displayed a more severe indole sensitivity than *iss1*-*1* alone.

**Figure 7.**
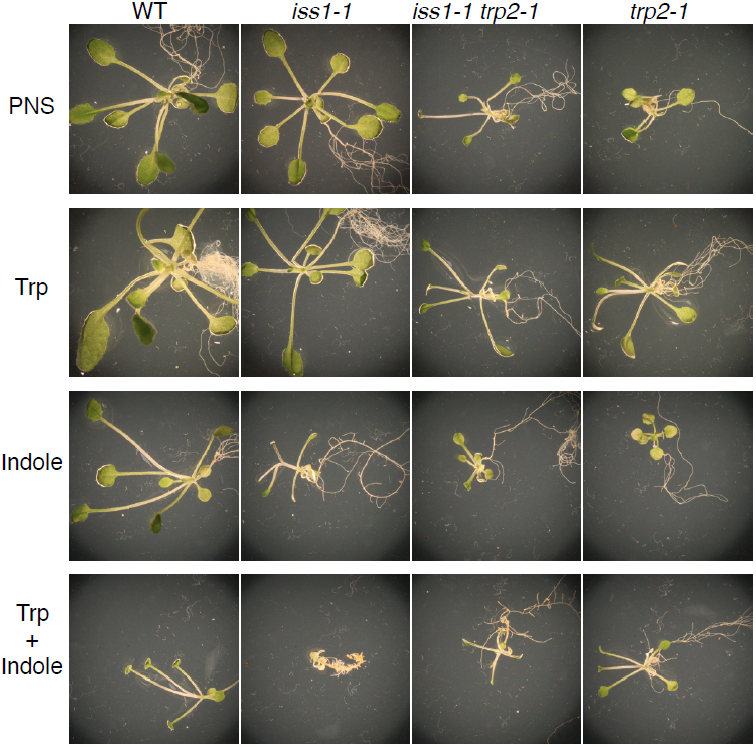
Analysis of the *iss1*-*1 trp2*-*1* double mutant phenotype. Shown are representative 21-day-old WT, *iss1*-*1, iss1*-*1 trp2*-*1* and *trp2*-*1* plants grown on PNS(top row), or PNS containing 80 μM Trp (second row), 80 μM indole (third row), or 80 μM Trp 80 μM indole (bottom row).

To confirm that elevated Trp levels are important for the *iss1*-*1* phenotype, we compared the severity of the *iss1*-*1* phenotype of plants grown on PNS media with either added indole, Trp, or indole together with Trp. *iss1*-*1* plants grown on PNS+indole+Trp medium had a much more pronounced high-IAA phenotype compared to *iss1*-*1* plants grown on PNS+indole medium, consistent with elevated Trp levels contributing to the *iss1*-*1* high IAA phenotype (Figure 7).

### *iss1*-*1* makes normal levels of indolic glucosinolates

The increased sensitivity of *iss1*-*1* to 5MT as well as the indole-dependent increase in Trp could be explained by reduction in catabolism of Trp into indole glucosinolates (IGs). IGs are an important sink for Trp in Arabidopsis, and mutants with reduced IGs display increased sensitivity to 5MT (Celenza *et al*. 2005; Bender and Celenza 2009). The *iss1*-*1* mutant had similar IG levels to WT whether grown on PNS or PNS+indole media, indicating that conversion of Trp to IGs is not altered in the *iss1*-*1* mutant. Interestingly both *iss1*-*1* and WT showed a 1.5 fold increase in total IG levels when grown on PNS+indole medium (Table S6).

### *ISS1* encodes an aminotransferase

Using map-based cloning (Lukowitz *et al*. 2000) the *ISS1* gene was identified as At1g80360 (Figure S1). Comparison between the *iss1*-*1* and Col-0 At1g80360 genomic sequences revealed a complex mutation beginning at base 335 of the coding sequence. This mutation consists of a 20 base deletion and a 51 base insertion of T-DNA that would result in a severely truncated protein when expressed (Figure S1). Sequencing of the *iss1*-*2* allele revealed a single point mutation that changes arginine 362 to a Trp residue in the predicted protein (Figure S1). Introduction of an expression construct with the At1g80360 cDNA driven by the CaMV 35S promoter rescued both the *iss1*-*1* and *iss1*-*2* mutants from the indole-dependent high IAA phenotype, confirming the identity of *ISS1* (Figure S2).

At1g80360 encodes a predicted fold-type I aminotransferase (AT), that was recently identified as VAS1, an AT capable of converting IPA to Trp (Zheng *et al*. 2013). ATs require pyridoxal 5’-phosphate (PLP) as a cofactor and catalyze the reversible reaction in which an amino group is transferred from an amino acid donor to a 2-oxo acid acceptor (Jensen and Gu 1996). BLAST analysis revealed that ISS1 is conserved across plant species; however, none of these presumed orthologs have a characterized enzyme activity. In the Arabidopsis genome, there are no ISS1 paralogs; however the proteins most similar to ISS1 are the bifunctional aspartate/prephenate AT encoded by *MEE17* (24% amino acid identity) and the Tyr AT TAT3 (23% amino acid sequence identity) (Figure S3). ISS1 shares only 12% amino acid sequence identity with TAA1, another Arabidopsis Trp AT (Figure S3).

### Heterologous expression of *ISS1* rescues the yeast Phe and Tyr auxotroph mutant *aro8 aro9*

Because *iss1*-*1* mutants have altered Trp metabolism and the ISS1 protein is most similar to two enzymes involved in aromatic amino acid biosynthesis, we hypothesized that ISS1 could function as a general AroAT. To test this hypothesis we asked if heterologous expression of *ISS1* in *aro8 aro9* mutant yeast could rescue the Phe and Tyr auxotrophy. In yeast, the AroATs encoded by *ARO8* and *ARO9* convert phenylpyruvate (PPY) to Phe and 4-hydorxyphenylpyruvate (4-HY) to Tyr (Iraqui *et al*. 1998; Urrestarazu *et al*. 1998). As seen in Figure 8A, galactose-induced *ISS1* expression was able to rescue the *aro8 aro9* Phe and Tyr auxotropy. The *iss1*-*2* mutant cDNA did not rescue the *aro8 aro9* mutant indicating that the point mutant can no longer function as an AroAT (Figure 8A). *TAA1* weakly rescued the *aro8 aro9* under galactose induction, consistent with its in vitro activity primarily using Trp as a substrate (Tao *et al*. 2008). In addition, *MEE17* failed to rescue the yeast mutant consistent with its reported activity with prephenate, but not with Phe or Tyr as substrates (Maeda *et al*. 2010) (Figure 8A).

**Figure 8.**
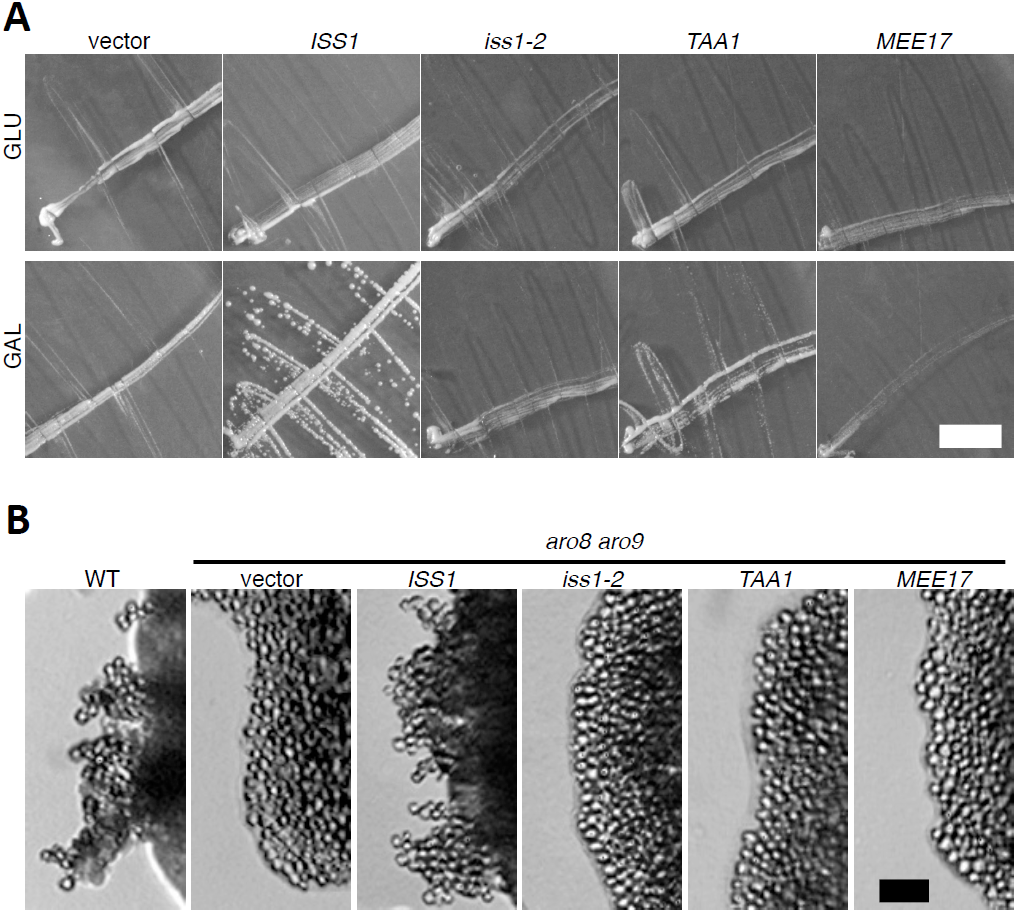
Heterologous expression of *ISS1* rescues the *aro8 aro9* mutant yeast. (A) pHY326-loxH (vector) or pHY326-loxH expressing *ISS1, iss1*-*2*,*TAA1* or *MEE17* were transformed into *aro8 aro9* double mutant haploid yeast (MPY1) in the S288C background and grown on SC medium –Phe and –Tyr containing glucose (top panels)or galactose (botto panels) for 48h at 30° White bar = 5 mm. (B) The same plasmidsused in (A) were transformed into *aro8 aro9* double mutant diploid yeast (MPY5) in the ∑1278b background and grown on SLAD medium with either glucose (not shown) or galactose for 72h at 30°.WT is strain PY652. Black bar = 20 mm

Many species of fungi, including certain backgrounds of *S.cerevisiae*, are able to transition from a vegetative state to filamentous state under conditions of low nitrogen and high cell density (Gimeno *et al*. 1992). High cell density and low nitrogen availability cause the increased production of tryptophol (Trp-OH) and phenylethanol (Phe-OH) by deamination of Trp and Phe, respectively, which in turn causes the transition from vegetative growth to filamentous growth in diploid yeast (Chen and Fink 2006). The *aro8 aro9* double mutant fails to produce Trp-OH or Phe-OH, and therefore is unable to transition to filamentous growth (Chen and Fink 2006). To test if *ISS1* also functions catabolically as an AroAT, we grew *aro8 aro9* yeast expressing *ISS1* on low nitrogen medium (SLAD) with galactose as carbon source and observed a weak rescue of the filamentous growth phenotype (Figure 8B), indicating that *ISS1* is able to participate both in the biosynthesis of Phe and Tyr and in the catabolism of Phe and/or Trp. *aro8 aro9* yeast expressing *iss1*-*2, TAA1* or *MEE17* failed to develop filaments (Figure 8B).

### Recombinant ISS1 protein has aromatic aminotransferase activity

Mutant alleles of *ISS1* called *vas1* (*re<u>v</u>ers<u>a</u>l of <u>s</u>av3 phenotype 1*) have been reported previously (Zheng *et al*. 2013). These authors determined that VAS1 uses IPA as the amino acceptor to make Trp as a counterbalance to the conversion of Trp to IPA by TAA1. Because the *iss1-1* mutant phenotype suggested a role in Trp catabolism and heterologous expression of ISS1 demonstrated a role in Phe/Tyr metabolism, we further characterized AroAT activity of heterologously expressed ISS1. The *ISS1* wild-type cDNA and the *iss1*-*2* mutant allele were cloned into the pMAL-c4X vector in order to express proteins with N-terminal maltose binding protein (MBP) tags. To confirm the fusion proteins were still active as AroATs, the fusion proteins were transformed into the *DL39 E. coli* mutant. *DL39* has mutations in *tyrB* (Tyr AT), *aspC* (aspartate AT) and *ilvE* (branched chain AT) and is auxotrophic for Tyr, Phe, aspartic acid, leucine, isoleucine and valine (Riewe *et al*. 2012). Expression of MBP-ISS1, but not MBP-iss1-2 rescued the DL39 Phe and Tyr auxotrophy confirming that the MBP tag was not hindering the activity of the recombinant enzymes and also confirming ISS1’s AroAT activity (data not shown).

Following purification using the MBP tag, we confirmed MBP-ISS1 was able to use Trp, Phe, and Tyr as substrates *in vitro*, consistent with ISS1 being an AroAT. In addition, the reverse reactions were also catalyzed by MBP-ISS1 (Table S7) indicating that ISS1 is an AroAT.

### The *iss1*-*1* mutant has an altered phenylpropanoid profile

Our analyses of the *iss1* mutant indicate a role for *ISS1* in Trp metabolism. However, based on the yeast heterologous expression experiments and the in vitro assays, ISS1 is a broader AroAT with activities not limited to only Trp metabolism. To determine if *iss1*-*1* mutant plants also display altered Phe metabolism we tested *iss1*-*1* for sensitivity to the toxic Phe analog PFP. Like 5MT, PFP is incorporated into proteins instead of Phe and feedback inhibits chorismate mutase (CM) (Palmer and Widholm 1975). We found that *iss1*-*1* plants were more resistant to PFP compared to WT suggesting that *iss1*-*1* plants may have relaxed feedback inhibition or increased Phe turnover (Figure 2D).

The fact that Phe levels are not elevated in the *iss1*-*1* mutant relative to WT (Figure 6B) suggests that feedback inhibition is not dramatically altered in *iss1*-*1* mutant plants. To determine if Phe turnover is disrupted in *iss1*-*1* mutant plants, we quantified phenylpropanoid levels. *iss1*-*1* plants grown on PNS medium had a significant decrease in coniferin, a monolignol glucoside, while flavonoid levels were similar to WT (Figure 9). Interestingly, in *iss1*-*1* plants grown on PNS+indole medium, coniferin levels were significantly higher compared to growth on PNS medium, but these levels were still were significantly lower than WT plants when grown on PNS+indole medium.

**Figure 9.**
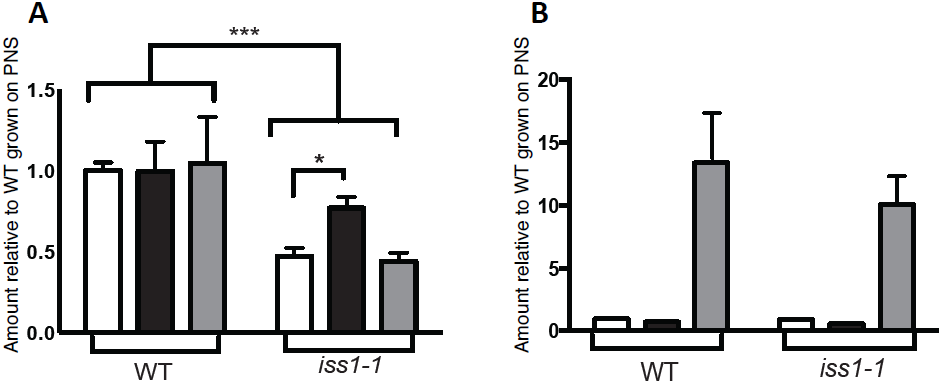
The *iss1* mutant has altered phenylpropanoid metabolism. Relative levels of coniferin (A) and flavonoids (B) are presented normalized to WT plants grown on PNS. Samples were extracted from whole 15-day-old Arabidopsis seedlings grown on PNS (white bars), PNS+indole (black bars), or PNS following 24 UV-B exposure (gray bars) and detected as described in Materials and Methods. For coniferin, upper brackets indicate a significantdecrease in coniferin in *iss1*-*1* compared to WT (***P< 0.0005 two-way ANOVA). Lower bracket indicates a significant increase in coniferin levels in iss1-1 when grown on indole compared to PNS (*P< 0.005 two-tailed Studen’s *t*-test). For flavonoids, a significant increase was found after UV-B treatment for both WT and *iss1*-*1* (P < 0.0001 two-way ANOVA).

Exposure to UV-B activates the production of certain phenylpropanoids, including some flavonoids (Ryan *et al*. 2001; Tohge *et al*. 2011). To determine whether the *iss1*-*1* mutant is affected in UV-B inducible flavonoid production, we measured phenylpropanoid levels after 24 h exposure to UV-B (Figure 9). After UV-B induction, several flavonoids increased approximately 12-fold in both WT and *iss1*-*1* plants while coniferin levels were unaffected by UV-B.

## Discussion

### The *iss1* mutant reveals increased use of Trp-independent IAA synthesis

Using a screen designed to find mutants with elevated indole-dependent IAA synthesis, two alleles of *ISS1* were identified that displayed a high IAA phenotype when grown on indole but not when grown on Trp. The *iss1*-*1* mutation is a complex insertion/deletion and *iss1*-*2* changes an amino acid conserved in other aminotransferases, thus we suspect that both alleles are null or severe hypomorphs. When grown on indole (but not Trp), both *iss1* mutant alleles display narrowed leaves and increased lateral and adventitious root growth, phenotypes consistent with elevated IAA levels.

In support of the indole-dependent growth phenotype, *iss1*-*1* seedlings grown on indole showed an eight-fold increase in IAA levels compared to WT grown under the same conditions; however IAA levels were similar to WT levels when grown on unsupplemented medium. We used dual labeling with [^15^N] ANA (an upstream precursor of Trp) and [^13^C_11_ ^15^N_2_] Trp to determine that these indole-dependent elevated IAA levels were due to an increase in Trp-I IAA biosynthesis. Our data demonstrate that on unsupplemented medium the majority of labeled IAA in both *iss1*-*1* and WT is from a Trp-D IAA pathway, consistent with what has been reported previously for wild-type Arabidopsis seedlings (Quint *et al*. 2009; Yu 2014). This finding suggests that Trp-D IAA biosynthesis is not obviously disrupted in *iss1*-*1* plants under normal lab growth conditions and is consistent with our finding that IAA levels are similar for WT and *iss1-1* grown without an indole supplement. In contrast, indole grown *iss1*-*1* seedlings had a higher percentage of [^15^N] IAA compared to [^15^N] Trp, consistent with increased use of Trp-I IAA synthesis, while WT plants grown on indole used primarily Trp-D IAA synthesis. These labeling results strongly suggest that the *iss1*-*1* mutant has an indole-dependent increase in Trp-I IAA synthesis. Both WT and *iss1*-*1* grown with or without an indole supplement showed nearly identical ratios of [^15^N] Trp to [^13^C_11_ ^15^N_2_] Trp incorporation into the Trp pool. This finding indicates that uptake of [^15^N] ANA is not perturbed in the *iss1*-*1* mutant and is consistent with indole-grown *iss1*-*1* having increased Trp-I IAA synthesis.

Consistent with this model, we found that the *iss1*-*1 taa1* double mutant and the *iss1*-*1 cyp79B2 cyp79B3* triple mutant showed the same degree of indole sensitivity as the single *iss1*-*1* mutant suggesting that the IPA and IAOx Trp-D IAA synthesis pathways are not responsible for the increased IAA synthesis in indole-grown *iss1*-*1* plants.

### *ISS1* functions in Trp catabolism

Surprisingly, *iss1*-*1* seedlings also have a 174-fold increase in Trp when grown on indole indicating that the increased usage of Trp-I IAA synthesis may be due indirectly to a loss of Trp catabolism. Consistent with decreased Trp catabolism, *iss1* has increased sensitivity to 5MT. In addition, we found that the indole-sensitive phenotype depends on having a functional Trp synthase b. The *trp2*-*1* mutation in *TSB1* suppresses the *iss1*-*1* indole-dependent growth phenotype consistent with a buildup of Trp being required for increased usage of Trp-I IAA synthesis. This suppression also indicates that the increased Trp-I IAA synthesis is not due simply to a build-up of indole and also suggests that Trp-I IAA synthesis branches off from IGP and not indole. Consistent with high Trp levels being responsible for the *iss1* phenotype, growth of the *iss1-1 trp2*-*1* double mutant on a combination of 80 μM Trp and 80 μM indole restored the *iss1*-*1* mutant phenotype (Figure 7). In addition, single *iss1*-*1* mutant seedlings grown under this condition showed a more severe phenotype than *iss1*-*1* mutants grown on 80 μM indole alone (Figure 7). *iss1*-*1* plants grown on 80 μM Trp appeared similar to WT grown in the same condition (Figure 7).

The Trp catabolism pathway that is affected in *iss1* mutants is undetermined at this time. In yeast and bacteria, Trp-derived IPA is converted to Trp-OH via the Ehrlich pathway (Hazelwood *et al*. 2008). While Trp-OH has been detected in cucumber seedlings (Rayle and Purves 1967) more recent studies with peas revealed very low levels of Trp-OH(Quittenden *et al*. 2009).

### Where does *ISS1* fit into IAA metabolism?

Mutant alleles of *ISS1*, called *vas1*, were identified as suppressors of the *taa1* mutant phenotype; failure of hypocotyl elongation in response to shade (Zheng *et al*. 2013). Under normal growth conditions, *vas1* mutants showed subtle increases in hypocotyl and petiole lengths and a significant increase in IAA and IPA levels suggesting that *VAS1* has altered IPA/YUC dependent IAA metabolism (Zheng *et al*. 2013). Consistent with a role for *VAS1* as a suppressor of TAA activity, the triple *vas1-2 sav3*-*1 tar2*-*1* mutant had normal hypocotyl length and is fertile (Zheng *et al*. 2013). Additionally, *vas1* mutant plants displayed a five-fold increase in the ethylene precursor ACC, pointing to a link between IAA synthesis and ethylene metabolism (Zheng *et al*. 2013).

Because the *vas1* phenotype is suggestive of a role in the IPA/YUC pathway, recombinant VAS1 enzyme characterization focused on using Trp as an amino donor or IPA as an amino acceptor. Using IPA as the amino acceptor, VAS1 had the highest catalytic (Kcat/Km) activity with methionine as the amino donor followed by Phe as the amino donor (Zheng *et al*. 2013). Based on the *vas1* phenotype and the enzyme kinetics of the recombinant VAS1 protein, it was proposed that *VAS1* normally functions as a counterbalance to *TAA1* dependent production of IPA while also impacting ethylene synthesis (Zheng *et al*. 2013). In this view, *VAS1* preferentially catalyzes the reverse reaction converting excess IPA to Trp, thus reducing the levels of free IPA.

Our results are distinct from those reported for *VAS1* and suggest that this gene has an additional role in Trp (and possibly Phe and/or Tyr) metabolism. We note *iss1’s* indole-sensitivity phenotype is strongest in plants greater than 18 days old while the *vas1* phenotype was characterized in seven-to nine-day old seedlings. In addition, the aerial portions of seven-day old *vas1* seedlings were found to have a two-fold increase in IAA (Zheng *et al*. 2013), whereas we found that three-week old *iss1* whole seedlings had no significant change in IAA levels unless supplemented with indole. We also note that while *VAS1’s* expression overlaps with the meristem-specific expression of *TAA1* and *YUC* genes, gene expression databases show that *ISS1/VAS1* is expressed at similar levels in meristematic and vegetative tissues whereas *TAA1* and *YUC* genes are expressed at much lower levels in vegetative tissues compared to meristematic tissues (www.weigelworld.org/resources/microarray/AtGenExpress/). We propose that the differing roles for *ISS1/VAS1* in Trp metabolism depend on the tissue type and/or developmental timing. In meristematic tissues, where the expression of *TAA1* (Stepanova *et al*. 2008; Yamada *et al*. 2009) and *YUC* genes (Cheng *et al*. 2007) is highest, *ISS1/VAS1* modulates levels of IPA by converting excess IPA into Trp (Zheng *et al*. 2013). In vegetative tissues, where IAA biosynthetic genes are not highly expressed, *ISS1/VAS1* functions in Trp degradation, by converting Trp into IPA leading into an uncharacterized Trp degradation pathway.

Based on its role in modulating Trp and IAA levels, we do not think that *ISS1/VAS1* is directly part of a Trp-I IAA biosynthesis pathway; rather we propose that the loss of *ISS1/VAS1* activity in Trp homeostasis increases Trp-I IAA synthesis. A number of reports indicate that Trp-I and Trp-D IAA synthesis are under exquisite developmental and environmental regulation (Michalczuk *et al*. 1992; Ljung *et al*. 2001; Epstein *et al*. 2002; Rapparini *et al*. 2002; Ribnicky *et al*. 2002; Yamada *et al*. 2009); perhaps one way of regulating Trp-I IAA synthesis is by modulating Trp metabolic flux.

### ISS1 is a member of a third class of plant aromatic aminotransferases

We found that heterologous expression of either *ISS1* or *TAA1* rescued the Phe and Tyr auxotrophies of yeast and *E. coli* mutants defective in AroAT activity. However, only *ISS1* expression was able to rescue the filamentous growth phenotype of the diploid *aro8 aro9* yeast mutant, indicating that ISS1 can carry out both aromatic amino acid catabolism and biosynthesis in vivo. In agreement with these in vivo findings, ISS1 has in vitro AT activity using Phe, Tyr or Trp as the amino donor or amino acceptor.

Two distinct families of ATs are known to participate in plant aromatic amino acid metabolism: the TAA family of Trp ATs, and Tyr ATs (TATS). TAA1, and the related TAR proteins are more than 50% identical, while TAA1 and ISS1 are only 12% identical consistent with ISS1 not being in the TAA1 family. The seven Arabidopsis TATs share between 38-51% sequence identity with each other (Prabhu and Hudson 2010) while TATs and ISS1 share only 23% sequence identity. Two TATs were able to rescue the *DL39 E. coli* Phe and Tyr auxotrophy (Prabhu and Hudson 2010; Riewe *et al*. 2012), and were found to have AroAT activity *in vitro* (Prabhu and Hudson 2010; Riewe *et al*. 2012). Additionally, the *tat5* mutant has an eleven-fold increase in Tyr levels and a decrease in Tyr secondary metabolites while showing no changes in Phe or Trp metabolism (Riewe *et al*. 2012). Based on the phenotype of the *iss1* mutant and limited similarity of ISS1 to TAT proteins, we suggest that ISS1 is a member of a plant AroAT family distinct from those previously described. An open question is whether ISS1 contributes to Phe and/or Tyr biosynthesis. While plants primarily convert prephenate to Phe/Tyr by transamination followed by dehydration/decarboxylation (Tzin and Galili 2010), recent findings indicate that plants possess enzyme activities capable of carrying out the dehydration/decarboxylation of prephenate first, followed by transamination to Phe or Tyr (Tzin *et al*. 2009; Yoo *et al*. 2013). ISS1’s ability to rescue the yeast *aro8 aro9* mutant combined with its in vitro activities is consistent with ISS1 also serving a role in Phe/Tyr metabolism.

### *iss1* mutants reveal an interaction between Trp and Phe metabolism

Given that several recent reports have demonstrated a high degree of coordination between Trp and Phe metabolism (Yamada *et al*. 2008; Tzin *et al*. 2009; Huang *et al*. 2010), it was not surprising that *iss1* plants have altered Phe metabolism as exhibited by reduced coniferin production and altered sensitivity to PFP. In plants, the majority of Phe is used for the production of the two classes of phenylpropanoids, monolignols and flavonoids (Huang *et al*. 2010). On unsupplemented medium, *iss1*-*1* plants displayed a decrease in coniferin while flavonoid levels were unchanged. This finding suggests that the *iss1*-*1* mutation disrupts regulation of specific steps downstream of the general phenylpropanoid pathway. Environmental stresses such as UV-B regulate the phenylpropanoid pathway as exemplified by increased production of certain flavonoids (Ryan *et al*. 2001; Tohge *et al*. 2011). We observed similar fold increases in flavonoid levels between *iss1*-*1* plants and WT plants grown on unsupplemented medium following 24h UV-B exposure suggesting that *iss1*-*1* affects constitutive phenylpropanoid production rather than inducible phenylpropanoid synthesis.

The fact that *iss1*-*1* is resistant to PFP suggests an alteration in Phe metabolism. In plants, resistance to the toxic Phe analog PFP is associated with a loss of feedback inhibition (Palmer and Widholm 1975) or an increased demand for Phe (Berlin and Widholm 1977). We found that Phe levels in the *iss1*-*1* mutant are similar to WT, suggesting that feedback inhibition is not changed. In addition, the decrease in phenylpropanoid content in *iss1*-*1* would suggest decreased Phe turnover. Because the *iss1* mutant’s effects on Phe metabolism are subtle, these phenotypes may be due to direct disruption of a Phe metabolic pathway that ISS1 has a minor role in or to indirect effects caused by dysregulation of Trp metabolism.

## Conclusions

From this work and the recent characterization of *vas1*, ISS1/VAS1 is an AroAT that has multiple roles in Trp metabolism. ISS1/VAS1 functions in meristematic tissue to modulate IAA levels by conversion of IPA to Trp, while in vegetative tissues it functions in Trp catabolism. *iss1* mutant plants have an indole-dependent increase in Trp-I IAA synthesis that results in a significant increase in IAA. Additionally, the fact that high levels of Trp appear to be required for the redirection of indole into Trp-I IAA synthesis points to an important role for Trp catabolism in IAA homeostasis. That *iss1* has altered Trp as well as Phe metabolism supports the idea of coordinated flux of metabolites through the Trp and Phe biosynthetic pathways. ISS1/VAS1-related proteins appear across the plant kingdom and form a distinct clade within the α I family of PLP-dependent enzymes. This suggests a conserved function of ISS1/VAS1-related proteins in plant metabolism, and future investigations of ISS1/VAS1 will be useful for revealing interactions between primary and secondary aromatic amino acid metabolism.

## Acknowledgments

We thank Reeta Prusty-Rao (Worcester Polytechnic Institute, Worcester, MA) for yeast strains Y652 and Y653. This work was supported by grants from the US National Science Foundation (MCB 0517506 to MCB0724970 to J.L.C.; MCB 0517420 and MCB 0725192 to J.N.; DBI 0552858 to J.R.) and funds from Boston University’s Undergraduate Research Opportunities Program (C.F., J.G., N.T. and C.W.).

## Supporting Information

Additional Supporting Information may be found in the online version of this article:

**Table S1.** Positional cloning primers that amplify microsatellite variations between Ws and Col-0 ecotypes.

**Table S2.** Plant genotyping oligonucleotides.

**Table S3.** Molecular and fragment ions of derivitized IAA and aromatic amino acids.

**Table S4.** Univector host plasmids and pUNI cDNAs used.

**Table S5.** *Saccharomyces cerevisiae* strains used.

**Table S6.** Indole glucosinolate quantification.

**Table S7.** Specific activity of purified MBP-ISS1.

**Figure S1.** Map-based cloning of the *ISS1* gene.

**Figure S2.** Overexpression of ISS1 cDNA rescues the *iss1* phenotype.

**Figure S3.** Amino acid sequence alignment of *ISS1* to fold type-I aminotransferases.

## Literature Cited

Ausubel, F. M., R. Brent, R. E. Kingston, D. D. Moore, J. G. Seidman et al., 1994-1998 Current Protocols in Molecular Biology. John Wiley and Sons, New York.

Bainbridge, K., S. Guyomarc’h, E. Bayer, R. Swarup, M. Bennett et al., 2008 Auxin influx carriers stabilize phyllotactic patterning. Genes Dev. 22: 810–823.

Barkawi, L. S., Y. Y. Tam, J. A. Tillman, J. Normanly and J. D. Cohen, 2010 A high-throughput method for the quantitative analysis of auxins. Nat. Protoc. 5: 1609–1618.

Bender, J., and J. L. Celenza, 2009 Indolic glucosinolates at the crossroads of tryptophan metabolism. Phytochem. Rev. 8: 25–37.

Berlin, J., and J. M. Widholm, 1977 Correlation between phenylalanine ammonia lyase activity and phenolic biosynthesis in p-fluorophenylalanine-sensitive and - resistant tobacco and carrot tissue cultures. Plant Physiol. 59: 550–553.

Brown, P. D., J. G. Tokuhisa, M. Reichelt and J. Gershenzon, 2003 Variation of glucosinolate accumulation among different organs and developmental stages of *Arabidopsis thaliana*. Phytochemistry 62: 471–481.

Celenza, J. L., Jr., P. L. Grisafi and G. R. Fink, 1995 A pathway for lateral root formation in *Arabidopsis thaliana*. Genes Dev. 9: 2131–2142.

Celenza, J. L., J. A. Quiel, G. A. Smolen, H. Merrikh, A. R. Silvestro et al., 2005 The Arabidopsis ATR1 Myb transcription factor controls indolic glucosinolate homeostasis. Plant Physiol. 137: 253–262.

Chen, H., and G. R. Fink, 2006 Feedback control of morphogenesis in fungi by aromatic alcohols. Genes Dev. 20: 1150–1161.

Chen, W. P., X. Y. Yang, A. D. Hegeman, W. M. Gray and J. D. Cohen, 2010 Microscale analysis of amino acids using gas chromatography-mass spectrometry after methyl chloroformate derivatization. J. Chromatogr. B Analyt. Technol. Biomed. Life Sci. 878: 2199–2208.

Cheng, Y., X. Dai and Y. Zhao, 2007 Auxin synthesized by the YUCCA flavin monooxygenases is essential for embryogenesis and leaf formation in Arabidopsis. Plant Cell 19: 2430–2439.

Clough, S. J., and A. F. Bent, 1998 Floral dip: a simplified method for *Agrobacterium-mediated* transformation of *Arabidopsis thaliana*. Plant J. 16: 735–743.

Epstein, E., J. Cohen and J. Slovin, 2002 The biosynthetic pathway for indole-3-acetic acid changes during tomato fruit development. Plant Growth Regul. 38: 15–20.

Gimeno, C. J., P. O. Ljungdahl, C. A. Styles and G. R. Fink, 1992 Unipolar cell divisions in the yeast S. cerevisiae lead to filamentous growth: regulation by starvation and RAS. Cell 68: 1077–1090.

Glawischnig, E., B. G. Hansen, C. E. Olsen and B. A. Halkier, 2004 Camalexin is synthesized from indole-3-acetaldoxime, a key branching point between primary and secondary metabolism in *Arabidopsis*. Proc. Natl. Acad. Sci. U S A 101: 8245–8250.

Guo, H., and J. R. Ecker, 2003 Plant responses to ethylene gas are mediated by SCF(EBF1/EBF2)-dependent proteolysis of EIN3 transcription factor. Cell 115: 667–677.

Haughn, G. W., and C. Somerville, 1986 Sulfonylurea-resistant mutants of *Arabidopsis thaliana*. Mol. Gen. Genet. 204: 430–434.

Hazelwood, L. A., J. M. Daran, A. J. van Maris, J. T. Pronk and J. R. Dickinson, 2008 The Ehrlich pathway for fusel alcohol production: a century of research on *Saccharomyces cerevisiae* metabolism. Appl. Environ. Microbiol. 74: 2259–2266.

Hemm, M. R., S. D. Rider, J. Ogas, D. J. Murry and C. Chapple, 2004 Light induces phenylpropanoid metabolism in Arabidopsis roots. Plant J. 38: 765–778.

Huang, T., T. Tohge, A. Lytovchenko, A. R. Fernie and G. Jander, 2010 Pleiotropic physiological consequences of feedback-insensitive phenylalanine biosynthesis in *Arabidopsis thaliana*. Plant J. 63: 823–835.

Hull, A. K., R. Vij and J. L. Celenza, 2000 Arabidopsis cytochrome P450s that catalyze the first step of tryptophan-dependent indole-3-acetic acid biosynthesis. Proc. Natl. Acad. Sci. U S A 97: 2379–2384.

Iraqui, I., S. Vissers, M. Cartiaux and A. Urrestarazu, 1998 Characterisation of Saccharomyces cerevisiae *ARO8* and *ARO9* genes encoding aromatic aminotransferases I and II reveals a new aminotransferase subfamily. Mol. Gen. Genet. 257: 238–248.

Jensen, R. A., and W. Gu, 1996 Evolutionary recruitment of biochemically specialized subdivisions of family I within the protein superfamily of aminotransferases. J. Bacteriol. 178: 2161–2171.

Kerhoas, L., D. Aouak, A. Cingoz, J. M. Routaboul, L. Lepiniec et al., 2006 Structural characterization of the major flavonoid glycosides from *Arabidopsis thaliana* seeds. J. Agric. Food Chem. 54: 6603–6612.

Kleine-Vehn, J., L. Langowski, J. Wisniewska, P. Dhonukshe, P. B. Brewer et al., 2008 Cellular and molecular requirements for polar PIN targeting and transcytosis in plants. Mol. Plant 1: 1056–1066.

Last, R. L., P. H. Bissinger, D. J. Mahoney, E. R. Radwanski and G. R. Fink, 1991 Tryptophan mutants in Arabidopsis: the consequences of duplicated tryptophan synthase beta genes. Plant Cell 3: 345–358.

Liu, Q., M. Z. Li, D. Leibham, D. Cortez and S. J. Elledge, 1998 The univector plasmid-fusion system, a method for rapid construction of recombinant DNA without restriction enzymes. Curr. Biol. 8: 1300–1309.

Liu, X., A. D. Hegeman, G. Gardner and J. D. Cohen, 2012 Protocol: High-throughput and quantitative assays of auxin and auxin precursors from minute tissue samples. Plant Methods 8: 31.

Ljung, K., 2013 Auxin metabolism and homeostasis during plant development. Development 140: 943–950.

Ljung, K., A. Ostin, L. Lioussanne and G. Sandberg, 2001 Developmental regulation of indole-3-acetic acid turnover in Scots pine seedlings. Plant Physiol. 125: 464–475.

Lukowitz, W., C. S. Gillmor and W. R. Scheible, 2000 Positional cloning in Arabidopsis. Why it feels good to have a genome initiative working for you. Plant Physiol. 123: 795–805.

Maeda, H., A. K. Shasany, J. Schnepp, I. Orlova, G. Taguchi et al., 2010 RNAi suppression of arogenate dehydratase1 reveals that phenylalanine is synthesized predominantly via the arogenate pathway in petunia petals. Plant Cell 22: 832–849.

Mano, Y., K. Nemoto, M. Suzuki, H. Seki, I. Fujii et al., 2010 The AMI1 gene family: indole-3-acetamide hydrolase functions in auxin biosynthesis in plants. J. Exp. Bot. 61: 25–32.

Mashiguchi, K., K. Tanaka, T. Sakai, S. Sugawara, H. Kawaide et al., 2011 The main auxin biosynthesis pathway in Arabidopsis. Proc. Natl. Acad. Sci. U S A 108: 18512–18517.

Michalczuk, L., T. J. Cooke and J. D. Cohen, 1992 Auxin levels at different stages of carrot somatic embryogenesis. Phytochemistry 31: 1097–1103.

Mikkelsen, M. D., P. Naur and B. A. Halkier, 2004 Arabidopsis mutants in the C-S lyase of glucosinolate biosynthesis establish a critical role for indole-3-acetaldoxime in auxin homeostasis. Plant J. 37: 770–777.

Morant, M., C. Ekstrom, P. Ulvskov, C. Kristensen, M. Rudemo et al., 2010 Metabolomic, transcriptional, hormonal, and signaling cross-talk in superroot2. Mol. Plant 3: 192–211.

Nonhebel, H., Y. Yuan, H. Al-Amier, M. Pieck, E. Akor et al., 2011 Redirection of tryptophan metabolism in tobacco by ectopic expression of an Arabidopsis indolic glucosinolate biosynthetic gene. Phytochemistry 72: 37–48.

Normanly, J., J. D. Cohen and G. R. Fink, 1993 *Arabidopsis thaliana* auxotrophs reveal a tryptophan-independent biosynthetic pathway for indole-3-acetic acid. Proc. Natl. Acad. Sci. U S A 90: 10355–10359.

Ouyang, J., X. Shao and J. Li, 2000 Indole-3-glycerol phosphate, a branchpoint of indole-3-acetic acid biosynthesis from the tryptophan biosynthetic pathway in *Arabidopsis thaliana*. Plant J. 24: 327–333.

Palmer, J. E., and J. Widholm, 1975 Characterization of carrot and tobacco cell cultures resistant to p-fluorophenylalanine. Plant Physiol. 56: 233–238.

Peret, B., B. De Rybel, I. Casimiro, E. Benkova, R. Swarup et al., 2009 Arabidopsis lateral root development: an emerging story. Trends Plant Sci. 14: 399–408.

Pollmann, S., D. Neu and E. W. Weiler, 2003 Molecular cloning and characterization of an amidase from *Arabidopsis thaliana* capable of converting indole-3-acetamide into the plant growth hormone, indole-3-acetic acid. Phytochemistry 62: 293–300.

Prabhu, P. R., and A. O. Hudson, 2010 Identification and partial characterization of an L-tyrosine aminotransferase (TAT) from *Arabidopsis thaliana*. Biochem. Res. Int. 2010: 549572.

Quint, M., L. S. Barkawi, K. T. Fan, J. D. Cohen and W. M. Gray, 2009 Arabidopsis IAR4 modulates auxin response by regulating auxin homeostasis. Plant Physiol. 150: 748–758.

Quittenden, L. J., N. W. Davies, J. A. Smith, P. P. Molesworth, N. D. Tivendale et al., 2009 Auxin biosynthesis in pea: characterization of the tryptamine pathway. Plant Physiol. 151: 1130–1138.

Rapparini, F., Y. Y. Tam, J. D. Cohen and J. P. Slovin, 2002 Indole-3-acetic acid metabolism in *Lemna gibba* undergoes dynamic changes in response to growth temperature. Plant Physiol. 128: 1410–1416.

Rayle, D. L., and W. K. Purves, 1967 Isolation and identification of indole-3-ethanol (tryptophol) from cucumber seedlings. Plant Physiol. 42: 520–524.

Ribnicky, D. M., J. D. Cohen, W. S. Hu and T. J. Cooke, 2002 An auxin surge following fertilization in carrots: a mechanism for regulating plant totipotency. Planta 214: 505–509.

Riewe, D., M. Koohi, J. Lisec, M. Pfeiffer, R. Lippmann et al., 2012 A tyrosine aminotransferase involved in tocopherol synthesis in Arabidopsis. Plant J. 71: 850–859.

Ryan, K. G., E. E. Swinny, C. Winefield and K. R. Markham, 2001 Flavonoids and UV photoprotection in Arabidopsis mutants. Z. Naturforsch. C 56: 745–754.

Stepanova, A. N., J. Robertson-Hoyt, J. Yun, L. M. Benavente, D. Y. Xie et al., 2008 TAA1-mediated auxin biosynthesis is essential for hormone crosstalk and plant development. Cell 133: 177–191.

Stepanova, A. N., J. Yun, L. M. Robles, O. Novak, W. He et al., 2011 The Arabidopsis YUCCA1 flavin monooxygenase functions in the indole-3-pyruvic acid branch of auxin biosynthesis. Plant Cell 23: 3961–3973.

Sugawara, S., S. Hishiyama, Y. Jikumaru, A. Hanada, T. Nishimura et al., 2009 Biochemical analyses of indole-3-acetaldoxime-dependent auxin biosynthesis in Arabidopsis. Proc. Natl. Acad. Sci. U S A 106: 5430–5435.

Tao, Y., J. L. Ferrer, K. Ljung, F. Pojer, F. X. Hong et al., 2008 Rapid synthesis of auxin via a new tryptophan-dependent pathway is required for shade avoidance in plants. Cell 133: 164–176.

Tohge, T., M. Kusano, A. Fukushima, K. Saito and A. R. Fernie, 2011 Transcriptional and metabolic programs following exposure of plants to UV-B irradiation. Plant Signal Behav. 6: 1987–1992.

Tzin, V., and G. Galili, 2010 The biosynthetic pathways for shikimate and aromatic amino acids in *Arabidopsis thaliana*. Arabidopsis Book 8: e0132.

Tzin, V., S. Malitsky, A. Aharoni and G. Galili, 2009 Expression of a bacterial bifunctional chorismate mutase/prephenate dehydratase modulates primary and secondary metabolism associated with aromatic amino acids in Arabidopsis. Plant J. 60: 156–167.

Urrestarazu, A., S. Vissers, I. Iraqui and M. Grenson, 1998 Phenylalanine-and tyrosine-auxotrophic mutants of *Saccharomyces cerevisiae* impaired in transamination. Mol. Gen. Genet. 257: 230–237.

Weijers, D., M. Sauer, O. Meurette, J. Friml, K. Ljung et al., 2005 Maintenance of embryonic auxin distribution for apical-basal patterning by PIN-FORMED-dependent auxin transport in Arabidopsis. Plant Cell 17: 2517–2526.

Won, C., X. Shen, K. Mashiguchi, Z. Zheng, X. Dai et al., 2011 Conversion of tryptophan to indole-3-acetic acid by TRYPTOPHAN AMINOTRANSFERASES OF ARABIDOPSIS and YUCCAs in Arabidopsis. Proc. Natl. Acad. Sci. U S A 108: 18518–18523.

Woodward, A. W., and B. Bartel, 2005 Auxin: regulation, action, and interaction. Ann. Bot. 95: 707–735.

Wright, A. D., M. B. Sampson, M. G. Neuffer, L. Michalczuk, J. P. Slovin et al., 1991 Indole-3-acetic acid biosynthesis in the mutant maize orange pericarp, a tryptophan auxotroph. Science 254: 998–1000.

Yamada, M., K. Greenham, M. J. Prigge, P. J. Jensen and M. Estelle, 2009 The TRANSPORT INHIBITOR RESPONSE2 gene is required for auxin synthesis and diverse aspects of plant development. Plant Physiol. 151: 168–179.

Yamada, T., F. Matsuda, K. Kasai, S. Fukuoka, K. Kitamura et al., 2008 Mutation of a rice gene encoding a phenylalanine biosynthetic enzyme results in accumulation of phenylalanine and tryptophan. Plant Cell 20: 1316–1329.

Yoo, H., J. R. Widhalm, Y. Qian, H. Maeda, B. R. Cooper et al., 2013 An alternative pathway contributes to phenylalanine biosynthesis in plants via a cytosolic tyrosine:phenylpyruvate aminotransferase. Nat. Commun. 4: 2833.

Yu, P., 2014 New analytical methodologies in the study of auxin biochemistry. Dissertation in Plant Biological Sciences. University of Minnesota.

Zhao, Y., 2012 Auxin biosynthesis: a simple two-step pathway converts tryptophan to indole-3-acetic acid in plants. Mol. Plant 5: 334–338.

Zhao, Y., 2014 Auxin biosynthesis. Arabidopsis Book 12: e0173.

Zhao, Y., A. K. Hull, N. R. Gupta, K. A. Goss, J. Alonso et al., 2002 Trp-dependent auxin biosynthesis in Arabidopsis: involvement of cytochrome P450s CYP79B2 and CYP79B3. Genes Dev. 16: 3100–3112.

Zheng, Z., Y. Guo, O. Novak, X. Dai, Y. Zhao et al., 2013 Coordination of auxin and ethylene biosynthesis by the aminotransferase VAS1. Nat. Chem. Biol. 9: 244–246.

Zhou, Z. Y., C. G. Zhang, L. Wu, C. G. Zhang, J. Chai et al., 2011 Functional characterization of the CKRC1/TAA1 gene and dissection of hormonal actions in the Arabidopsis root. Plant J. 66: 516–527.

